# Prospects for genomic selection in cassava breeding

**DOI:** 10.1101/108662

**Authors:** Marnin D. Wolfe, Dunia Pino Del Carpio, Olumide Alabi, Chiedozie Egesi, Lydia C. Ezenwaka, Ugochukwu N. Ikeogu, Robert S. Kawuki, Ismail S. Kayondo, Peter Kulakow, Roberto Lozano, Ismail Y. Rabbi, Esuma Williams, Alfred A. Ozimati, Jean-Luc Jannink

## Abstract

Cassava (*Manihot esculenta* Crantz) is a clonally propagated staple food crop in the tropics. Genomic selection (GS) reduces selection cycle times by the prediction of breeding value for selection of unevaluated lines based on genome-wide marker data. GS has been implemented at three breeding programs in sub-Saharan Africa. Initial studies provided promising estimates of predictive abilities in single populations using standard prediction models and scenarios. In the present study we expand on previous analyses by assessing the accuracy of seven prediction models for seven traits in three prediction scenarios: (1) cross-validation within each population, (2) cross-population prediction and (3) cross-generation prediction. We also evaluated the impact of increasing training population size by phenotyping progenies selected either at random or using a genetic algorithm. Cross-validation results were mostly consistent across breeding programs, with non-additive models like RKHS predicting an average of 10% more accurately. Accuracy was generally associated with heritability. Cross-population prediction accuracy was generally low (mean 0.18 across traits and models) but prediction of cassava mosaic disease severity increased up to 57% in one Nigerian population, when combining data from another related population. Accuracy across-generation was poorer than within (cross-validation) as expected, but indicated that accuracy should be sufficient for rapid-cycling GS on several traits. Selection of prediction model made some difference across generations, but increasing training population (TP) size was more important. In some cases, using a genetic algorithm, selecting one third of progeny could achieve accuracy equivalent to phenotyping all progeny. Based on the datasets analyzed in this study, it was apparent that the size of a training population (TP) has a significant impact on prediction accuracy for most traits. We are still in the early stages of GS in this crop, but results are promising, at least for some traits. The TPs need to continue to grow and quality phenotyping is more critical than ever. General guidelines for successful GS are emerging. Phenotyping can be done on fewer individuals, cleverly selected, making for trials that are more focused on the quality of the data collected.

**Abbreviations:**

(GS): Genomic selection
(GBS): genotype-by-sequencing
(IITA): International Institute of Tropical Agriculture
(NRCRI): National Root Crops Research Institute
(NaCRRI): National Crops Resources Research Institute
(GEBVs): genomic estimated breeding values
(TP): training population
(RTWT): fresh root weight
(RTNO): root number
(SHTWT): fresh shoot weight
(HI): harvest index
(DM): dry matter
(CMD): content cassava mosaic disease
(MCMDS): mean CMD severity
(VIGOR): early vigor

## INTRODUCTION

Cassava (*Manihot esculenta* Crantz), a root crop with origins in the Amazon basin (Olsen and Schaal, 1999), provides staple food for more than 500 million people worldwide (Howeler et al., 2013). It is widely cultivated in Sub-Saharan Africa where the storage roots serve as primary source of carbohydrates and can be processed into a wide variety of products such as Fufu, Lafun, Gari, Abacha, Tapioca and starch (Chukwuemeka, 2007; Bamidele et al., 2015).

Cassava is a diploid (2n=36) and highly heterozygous non-inbred crop that is propagated vegetatively by farmers using stem cuttings, though most genotypes do flower and can be used to produce botanical seeds from either self or cross-pollination. Among the most important traits targeted for improvement are storage root yield, dry matter content, starch content, tolerance to postharvest physiological deterioration, carotenoids content and resistance to pests/diseases (Esuma et al., 2016).

Development and implementation of breeding strategies in cassava represent a challenge due to the crop’s heterozygous nature and long breeding cycle. A traditional cassava-breeding program relies on phenotypic characterization of mature plants that have been clonally propagated. Typically, cycles of selection take three to six years from seedling germination to multi-location yield trials and additional years are required for evaluation of promising genotypes before variety release (Figure 1).

**Figure 1.**
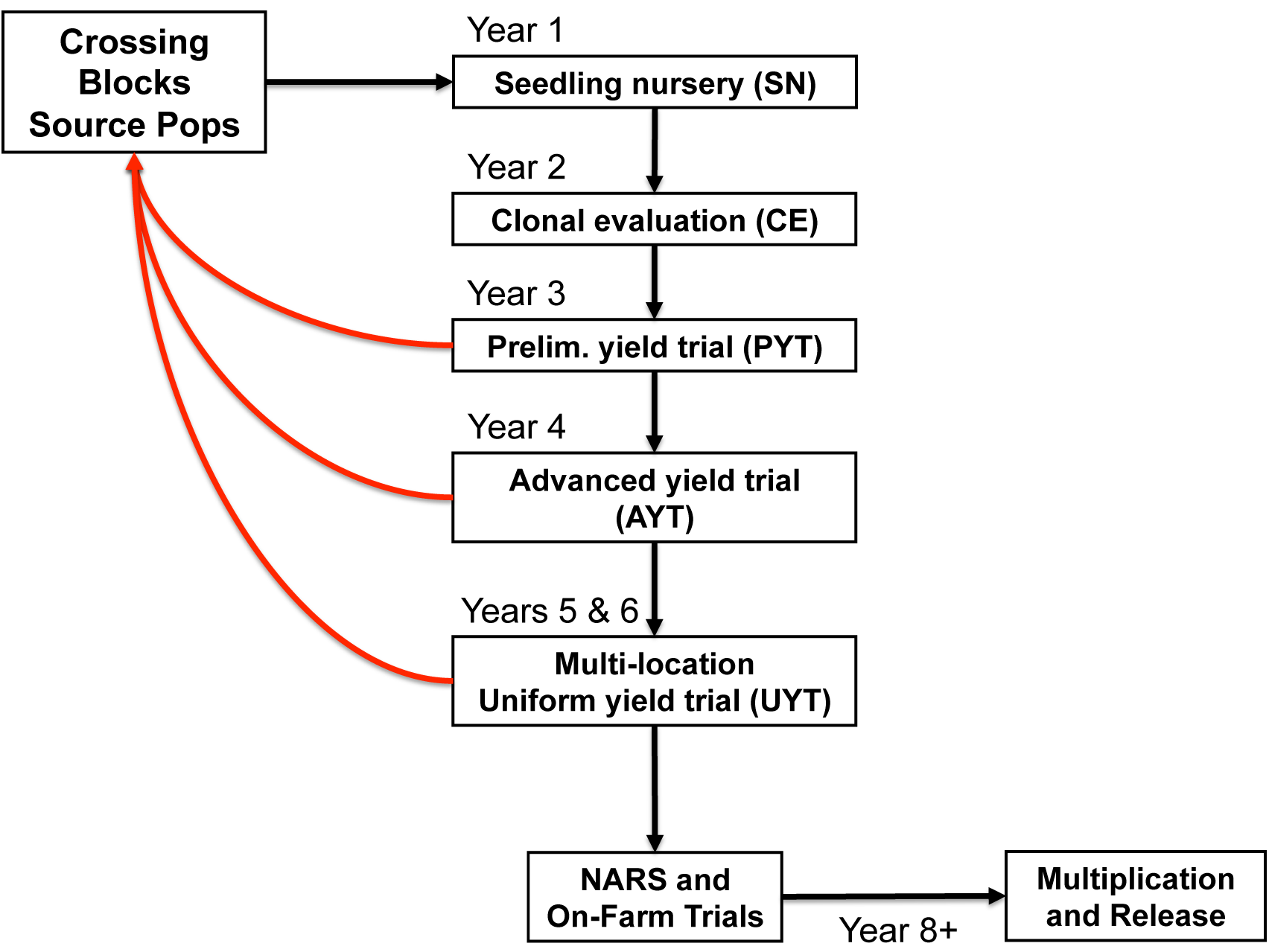
Schematic of a conventional cassava breeding cycle. Arrows between trials indicate the selection of materials for further phenotyping trials. Red arrows indicate the selection of materials as parents for crossing.

Marker-assisted selection (MAS) has been effective in cassava for the selection of promising genotypes for resistance to cassava mosaic disease (Okogbenin et al., 2007; Ceballos et al., 2015; Parkes et al., 2015). However, the use of MAS is limited primarily to monogenic traits, which makes this method infeasible for complex traits (Dekkers and Hospital, 2002; Heffner et al., 2009a).

With the advent of next generation sequencing technologies, it is now affordable to profile single nucleotide polymorphic (SNP) markers genome-wide (Barabaschi et al., 2015), which can support the use of genomic selection (GS), a breeding method that uses such markers to predict breeding values of unevaluated individuals (Meuwissen et al., 2001). GS can optimize and accelerate pipelines for population improvement, variety development and release (Heffner et al., 2009b) with reduction in breeding time due to selection of parental genotypes with superior breeding values at seedling stage based on genotypes alone.

In general, GS models differ with respect to the assumptions they make about genetic architecture. While random-regression (RRBLUP) and genomic-BLUP (GBLUP) models assume an infinitesimal genetic architecture (nearly equal and small contribution of all genomic regions to the phenotype), Bayesian methods are available that alter that assumption (Gianola et al., 2009; Legarra et al., 2011; Habier et al., 2011). Evaluation of different GS models using non-simulated data indicates that prediction accuracy varies across species and traits (Heslot et al., 2012; Resende et al., 2012; Gouy et al., 2013; Charmet et al., 2014; Rutkoski et al., 2014; Cros et al., 2015).

Previous studies in cassava have estimated genetic parameters and evaluated prediction accuracy applying the GBLUP model with small training sets and low-density markers (Oliveira et al., 2012, 2014). Historical phenotypic data from the International Institute of Tropical Agriculture (IITA) combined with markers obtained from genotyping-by-sequencing (GBS) showed promising results for cassava breeding using genomic selection (Ly et al., 2013). In that study, the predictive ability (accuracy) measured as the correlation between predictive values and the phenotypic value ranged from 0.15 to 0.47 across traits (Ly et al., 2013).

There are ongoing efforts under the Next Generation Cassava Breeding (NextGen Cassava) project (www.nextgencassava.org) to increase the rate of genetic improvement in cassava and unlock the full potential of cassava production. The project is currently in the early stages of implementing genomic selection at three African research institutes: the National Crops Resources Research Institute (NaCRRI) in Uganda, the National Root Crops Research Institute (NRCRI) and the IITA, both in Nigeria.

In the present study, we evaluated the potential of genomic selection as a breeding tool to increase rates of genetic gain in datasets associated with all three NextGen Cassava breeding programs. We assessed predictive ability by cross-validation within training population datasets for seven traits: dry matter (DM) content, fresh root weight (RTWT), root number (RTNO), shoot weight (SHTWT), harvest index (HI), severity of cassava mosaic disease (MCMDS) and plant vigor (VIGOR). We compared the performance of seven GS models for these traits in each of the breeding programs.

One important topic in genomic selection concerns the feasibility of prediction across generations and across training populations from different breeding populations or programs. To maximize the rate of gain achievable by GS, prediction models will need to accurately rank unevaluated progenies rather than genotypes contemporary to the training population. It is well known that recombination and divergence associated with recurrent selection reduces the accuracy of across-generation prediction, making this kind of prediction a major challenge for genomic selection. Accuracies in these scenarios have not been previously estimated in cassava. Therefore, we tested accuracy of across-generation prediction using the IITA training population and two successive cycles of progenies that have been phenotyped. Similarly, given that previous results indicated only a small genetic differentiation among clones from different populations (Wolfe et al., 2016a), we tested whether combining information from different populations could increase prediction accuracy in the smaller populations.

Finally, in a typical scenario a GS program will phenotype all selected materials and a subset of the unselected material in order to update the training model. We further investigated the impact of phenotyping different size subsets of materials for TP update. We compared random subset selections to selections based on a training population optimization algorithm (Akdemir et al. 2015).

This study is a starting point for successful application of genomic selection in African cassava. Similar to other studies, factors such as trait heritability, prediction model and training population composition play an important role. For example, traits with higher heritability like DM are considered to be more likely to respond to selection and lead to larger genetic gain over cycles of selection (Kawano et al 1998, Ceballos et al 2015). Our results will serve to guide implementation strategies for GS in cassava breeding programs.

## MATERIALS & METHODS

### Germplasm

In this study, we analyzed data from the genomic selection programs at three African cassava breeding institutions: NaCRRI, NRCRI and IITA. Germplasm from NaCRRI included 411 clones descended from crosses among accessions from East Africa, West Africa and South America. The collection from NRCRI was made up of 899 clones, 211 of them being in common with the IITA breeding germplasm. The remaining 688 clones were materials derived either in part or directly from the International Center for Tropical Agriculture (CIAT) in Cali, Columbia. Wolfe et al. (2016a) shows details of origins and pedigrees of the NaCRRI and NRCRI clones used in this study.

The primary IITA germplasm we have analyzed is also known as the Genetic Gain (GG) collection, which comprises 709 elite and historically important breeding clones and a few landraces that have been collected starting in the 1970’s. These materials have also been previously described in Okechukwu and Dixon (2008), Ly et al. (2013) and Wolfe et al. (2016a).

In addition, two generations of GS progenies were analyzed (Figure 2). The first, GS cycle 1 (C1) comprised 2,890 clones, from 166 full-sib (FS) families with 85 parents from the GG collection. Because successful crossing is a challenge in cassava, and in order to obtain the full set of desired matings among parents of C1, crossing blocks were planted in two successive years (2013 and 2014). In 2013, 79 parents produced 2,322 seedlings (135 FS families). In 2014, 17 parents, of which, 11 were re-used from the previous year and six were new parents from the GG collection, gave rise to an additional 568 seedlings (31 new FS families). C1 families have a mean size of 17.4 siblings (median 15, range 2 to 78).

**Figure 2.**
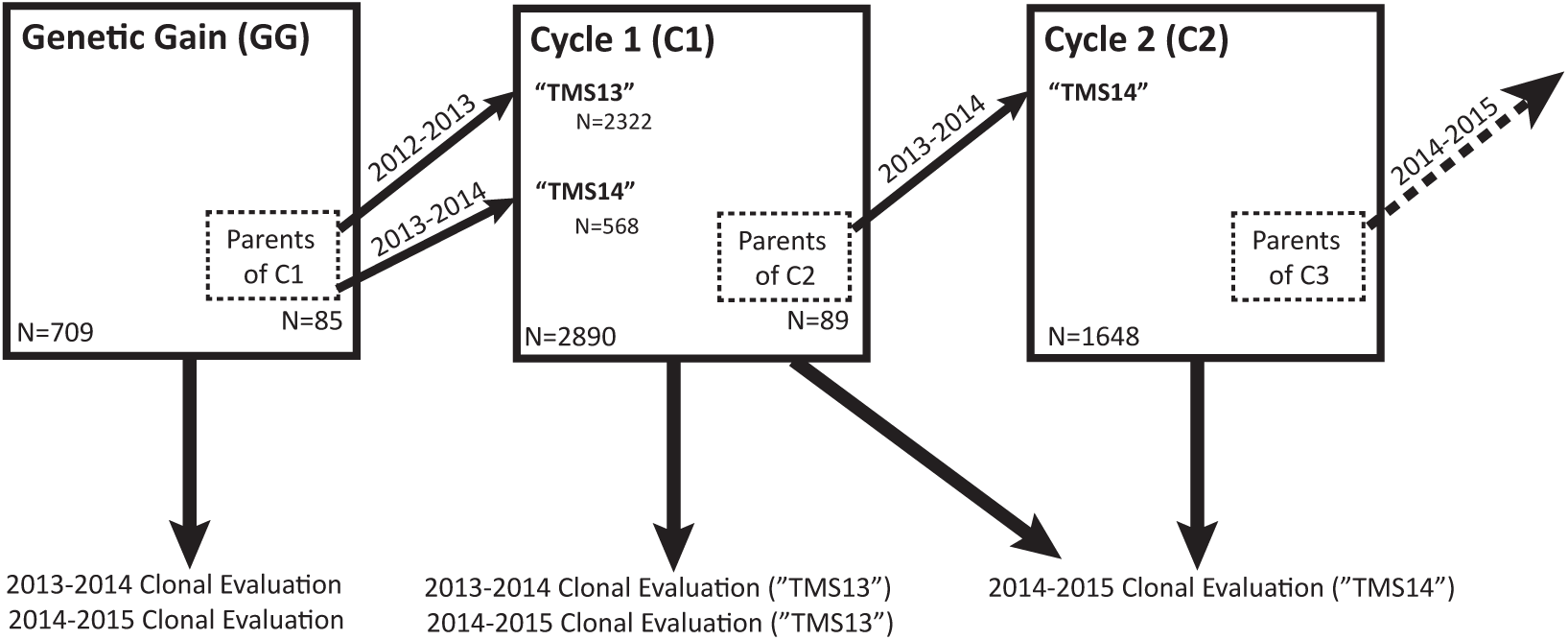
Schematic of IITA Genomic Selection 2012-2015. Three generations of IITA genomic selection program are illustrated here. From the genetic gain (GG) population, 85 parents were selected and crosses over two years (“TMS13F” in 2012-2013 and “TMS14F” in 2013-2014) gave rise to 2890 Cycle 1 (C1) progeny. Predictions based on data from the GG were used to select 89 parents from among C1 in 2013, giving rise to 1648 Cycle 2 (C2) progeny in 2014. The GG have been clonally evaluated in 2013-2014 and 2014-2015. The “TMS13” C1 were evaluated in 2013-2014 and 2014-2015. The “TMS14” C1 were evaluated with the C2 in 2014-2015.

Finally, in 2014, a crossing block was planted with 89 selected C1 parents and generated 1648 GS cycle 2 (C2) seedlings in 242 FS families. Cycle 2 families had a mean size of 6.8 individuals (median 6, range 1 to 20).

### Phenotyped traits

In total, seven traits were analyzed in this study. Plant vigor (VIGOR) was recorded as 3 (low), 5 (medium) and 7 (high), one month after planting (1 MAP) at IITA and NRCRI and three MAP at NaCRRI. We used the across-season average cassava mosaic disease severity score (MCMDS) for our analyses. MCMDS is the mean of measurements taken at 1, 3 and 6 MAP, on a scale of 1 (no symptoms) to 5 (severe symptoms). DM was expressed as a percentage of dry root weight relative to fresh root weight (RTWT). At IITA, DM was measured by drying 100 g of fresh roots in an oven whereas at NRCRI and NaCRRI, the specific gravity method (Kawano et al., 1987) was used. RTWT and SHTWT were expressed in kilograms per plot, whereas HI was the proportion of total biomass per plot that is RTWT. Meanwhile, RTNO was the number of fresh roots harvested per plot.

The phenotyping trials analyzed in this study have been described in part in previous publications (Wolfe et al. 2016a; Wolfe et al. 2016b). However, complete details on the phenotyping trial design particular to this study are provided in **Supplementary Methods**. All phenotyping trials were conducted between 2013 and 2015. NaCRRI clones were evaluated in three locations with different agro-ecological conditions in Uganda: Namulonge, Kasese and Ngetta. NRCRI clones were tested in three locations in Nigeria: Kano, Otobi and Umudike. Meanwhile, IITA clones were evaluated in four locations within Nigeria: Ibadan, Ikenne, Ubiaja and Mokwa.

### Two-stage genomic analyses

Except where noted, a two-step approach was used to evaluate genomic prediction in this study. This approach was used to correct for the heterogeneity in experimental designs and increase computational efficiency. The first stage involved accounting for trial-design related variables using a linear mixed model.

For NaCRRI we fitted the model: ***y = Xβ + Z*_*clone*_*c + Z*_*range(loc.year)*_*r + Z*_*block(range)*_*b + ɛ***, where ***β*** included a fixed effect for the population mean, the location-year combination and for plot-basis traits (RTWT, RTNO and SHTWT), the number of plants harvested per plot was included as a covariate; vector c and corresponding incidence matrix **Z_clone_** represented a random effect for clone where 
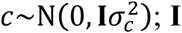
 represented the identity matrix, while the range variable was nested in location-year-replication and was represented by the incidence matrix **Z**_**range(loc.year)**_ and random effects vector 
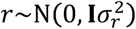
. Ranges were equivalent to a row or column along which plots were arrayed. Blocks were also modeled, with a block being a subset of a range. Block effects were nested in ranges and were incorporated as random with incidence matrix **Z_block(range)_** effects vector 
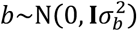
. Finally, the residuals ɛ were random, with 
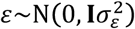
.

The model for NRCRI was: ***y*** = **X***β* + **Z_clone_***C* + **Z**_**set**(**loc.year**)_*S* + **Z**_**rep**(**set**)_*r* + **Z**_**block**(**rep**)_*b* + *ε*. Here, **Z_set_** was the incidence matrix corresponding to the random effect for the planting group (see above), which was nested in location-year, with 
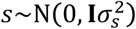
. Replication effects were nested in sets and treated as random with incidence matrix **Z_rep(set)_** and effects vector 
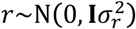
. Blocks were nested in replications, treated as random and represented by design matrix **Z_block(rep)_** and effects vector 
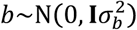
. The fixed effects for NRCRI included were the same as for NaCRRI, with the addition of a term for trial (i.e. TP1 and TP2; see above).

For IITA, data from all trials described above were fitted together using the following model: ***y*** = **X***β* + **Z_clone_***C* + **Z**_**range**(**loc.year**)_*r* + *ε*. The range effect was fit as random. The fixed effects were the same as those described for NaCRRI except the proportion of harvested plants (out of the total originally planted) was used instead of the number harvested as a cofactor. This was done to correct for differences in plot sizes.

BLUP (***ĉ***) for the clone effect, which represents an estimate of the total genetic value (EGV) for each individual, was extracted. EGVs were de-regressed by dividing by their reliability 
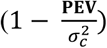
, where PEV is the prediction error variances of the BLUP. The mixed models above were solved using the *lmer* function of *lme4* package (Bates et al., 2014) in R.

For downstream genomic evaluations, we used the de-regressed EGVs and weighted error variances according to Garrick et al. (2009), using one divided by the square root of 
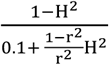
, where H^2^ is the proportion of the total variance explained by the clonal variance component, 
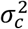
.

### Genotyping data

Cassava collections described above were genotyped using GBS (Elshire et al. 2011) with the *ApeKI* restriction enzyme recommended by Hamblin and Rabbi (2014). SNPs were called using the TASSEL 5.0 GBS pipeline v2 (Glaubitz et al., 2014) and aligned to cassava reference genome, v6.1 (http://phytozome.jgi.doe.gov; ICGMC, 2015). Genotype calls were only allowed when a minimum of two reads were present, otherwise the genotype was imputed (see below). Furthermore, the GBS data was filtered such that clones with >80% missing and markers with >60% missing genotype calls were removed. Markers with extreme deviation from Hardy-Weinberg equilibrium (Χ^2^ > 20) were also removed. Only biallelic SNP markers were considered for further analyses. We used a combination of custom scripts and common variant call file (VCF) (Danecek et al., 2011) manipulation tools to accomplish the above pipeline. Finally, imputation was conducted with Beagle v4.0 (Browning & Browning, 2009). A total of 155,871 markers were obtained following the procedures described above. For genomic prediction in a given population/dataset, we further filtered out SNPs with a minor allele frequency (MAF) less than 0.01.

### Assessment of prediction accuracy by cross-validation

In order to obtain unbiased estimates of prediction accuracy, we used a *k*-fold cross validation scheme (Kohavi, 1995). In brief, each breeding program dataset (NR, UG and GG) was split randomly into *k* = 5 fold mutually exclusive training and validation sets. The training set composed by four out of five of the folds was used to estimate marker effects for predictions. The estimated marker effects were used to predict the breeding value of validation set individuals. The process of fold assignment and genomic prediction was repeated 25 times for each model. For each repeat, predictions were accumulated from each individual when it was in the validation fold. Prediction accuracy was then calculated as the Pearson correlation (*cor* function in R) between the EGV and the accumulated predicted values for that repeat.

### Genomic prediction methods

In this study, we compared the accuracy of genomic prediction using seven methods that are briefly described below. These methods differ in their assumptions about genetic architecture and whether the prediction being made represents a genome estimated breeding value (GEBV, which includes additive effects) or a genome estimated total genetic value (GETGV, which includes additive plus non-additive effects). Prediction models were compared using several prediction scenarios (described in detail below), including 25 replications of 5-fold cross-validation, cross-generation and cross-population prediction.

**GBLUP**. Prediction with genomic BLUP (GBLUP) involves fitting a linear mixed model of the following form: ***y* = X*β*** + **Z*g*** + ***ɛ***. Here, ***y*** is a vector of the phenotype, ***β*** is a vector of fixed, non-genetic effects with design matrix **X**. The vector ***g*** is a random effect, the best linear unbiased prediction (BLUP), which represents the GEBV for each individual. **Z** is a design matrix pointing observations to genotype identities and ***ɛ*** is a vector of residuals. The GEBV is obtained by assuming 
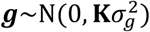
, where 
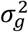
, is the additive genetic variance and **K** is the square, symmetric genomic realized relation matrix based on SNP marker dosages. The genomic relationship matrix used was constructed using the function *A.mat* in the R package rrBLUP (Endelman, 2011) and follows the formula of VanRaden (2008), method two. GBLUP predictions were made with the function *emmreml* in the R package EMMREML (Akdemir and Okeke, 2015).

**RKHS**. We made predictions using reproducing kernel Hilbert spaces (RKHS). The genomic relationship matrix used in the GBLUP model described above can be considered as a parametric (additive genetic) kernel function and exists as a special case of RKHS (Gianola and van Kaam, 2008; Morota and Gianola, 2014). For RKHS predictions, we used a mixed model of the same form as for GBLUP above. Unlike for GBLUP, we used a Gaussian kernel function: *K_ij_* = exp (−(d_ij_θ)). Here, K_ij_ was the measured relationship between two individuals, d_ij_ was their euclidean genetic distance based on marker dosages and θ was a tuning (sometimes called a “bandwidth”) parameter that determines the rate of decay of correlation among individuals. Because this is a nonlinear function, the kernels we used for RKHS could capture non-additive as well as additive genetic variation. Thus, the BLUPs from RKHS models represent GETGVs rather than GEBVs.

Because the optimal θ must be determined, a range of values was tested in two ways. First, we did cross-validation with the following θ values and selected the one with the best accuracy: 0.0000005, 0.000005, 0.00005, 0.0001, 0.0005, 0.001, 0.004, 0.006, 0.008, 0.01, 0.02, 0.04, 0.06, 0.08, 0.1 (Single kernel RKHS). Second, we used the *emmremlMultiKernel* function in the EMMREML package to fit a multiple-kernel model with six covariance matrices, with the following bandwidth parameters and allowed REML to find optimal weights for each: 0.0000005, 0.00005, 0.0005, 0.005, 0.01, 0.05 (Multi-kernel RKHS).

**Bayesian Marker Regressions**. We tested four well-established Bayesian prediction models: BayesCpi (Habier et al., 2011), the Bayesian LASSO (BL; Park and Casella, 2008), BayesA, and BayesB (Meuwissen et al., 2001). In ridge-regression (equivalent to GBLUP), marker effects are all shrunken by the same amount, because we assume they are all drawn from a normal distribution with the same variance. Further, all markers have nonzero effect and most have small effects, essentially assuming that the genetic architecture of the trait is infinitesimal. In contrast, the Bayesian models we tested allow for alternative genetic architectures by inducing differential shrinkage of marker effects. For BayesA and BL, all markers have nonzero effect but marker variances are drawn from scaled-t and double-exponential distributions respectively, which are both distributions with thicker tails and greater density at zero. BayesB and BayesCpi are variable selection models, because the marker variances come from a two-component mixture of a point mass at zero and either a scaled-t distribution (BayesB) or a normal distribution (BayesCpi). Fitting BayesB and BayesCpi begins by estimating a parameter *pi*, the proportion of markers with nonzero effect. We performed Bayesian predictions with the R package BGLR (Pérez and De Los Campos, 2014). Following Heslot et al. (2012) and others, we ran BGLR for 10,000 iterations, discarded the first 1000 iterations as burn-in and thinned to every 5^th^ sample. Marker dosages were mean-centered for each training population before analysis. Convergence was confirmed visually in initial test runs using the *coda* package in R (Plummer et al., 2006).

**Random Forest**. Random forest (RF) is a machine learning method used widely in regression and classification (Breiman, 2001; Strobl et al., 2009). The use of RF regression with marker data has been shown to capture epistatic effects and has been successfully used for prediction of GETGV (Breiman, 2001; Motsinger-Reif et al., 2008; Michaelson et al., 2010; Heslot et al., 2012; Charmet et al., 2014; Sarkar et al., 2015; Spindel et al., 2015). In prediction, a random forest is a collection of *r* regression trees grown on a subset of the original dataset that is bootstrapped over observations and randomly sampled over predictors. Averaging the prediction over trees for validation observations then aggregates information. We used RF with the parameter*, ntree* set to 500 and the number of variables sampled at each split (*mtry*) equal to 300. We implemented RF using the randomForest package in R (Liaw and Wiener, 2002). As in the Bayesian regressions, marker dosages were mean-centered before RF analysis.

### Comparison of models based on similarity of rankings

In order to test for GS model similarities among breeding programs we clustered the GEBV output on a breeding program basis. GEBVs from each model were scaled and centered on a column basis, using the *scale* function in R, and were then used to construct a matrix of Euclidean distances between models. Distance matrices were used as an input for hierarchical clustering using the Ward criterion implemented in the *hclust* R function (Heslot et al., 2012).

### Across-generation genomic predictions

Because nearly all of the IITA germplasm from C1 and C2 had been clonally evaluated, we were able to test the prospects for prediction of unevaluated progeny. We predicted all traits using all methods in four scenarios: GG predicts C1, GG predicts C2, C1 predicts C2, GG+C1 predicts C2. Unlike in the other predictions presented in this study, cross-generation predictions were done in a single step (raw phenotype and genomic data fit simultaneously). The exception was for RF, where correction for location and blocking factors is not supported. For RF prediction, we used the same de-regressed EGVs as for cross-validation. The software and parameters used were the same as already described. The design model is the same as described for IITA above.

### Training population update

We evaluated the impact on cross-generation prediction accuracy of phenotyping different size subsets of the un-selected C1 (materials selected for crossing in each cycle were phenotyped, but unselected materials were not phenotyped in all cases). We selected subsets of C1 using two methods: randomly and with a genetic algorithm implemented in the R package STPGA (Akdemir et al., 2015).

STPGA uses an approximation of the mean prediction error variance (PEV) expected for a given set of training individuals in combination with a given set of test genotypes as a criterion (which does not require phenotype data) for selecting the “optimal” training set. The genetic algorithm implemented by STPGA is used to rapidly find the training set that minimized the selection criterion (mean PEV of the test set; Akdemir et al., 2015). In order to speed up computation, STPGA uses principal components rather than raw SNP markers as genetic predictors.

Parents selected for further recombination were cloned into a crossing block. This is the point at which additional, un-selected seedlings must be chosen for phenotyping in order to incorporate their data in the prediction of the eventual progeny that are produced. Since the next generation of progenies had not yet been produced, we targeted STPGA on the parents of C2 (PofC2). Figure 3 provides a schematic of genomic selection with training population update and optimization using STPGA. We constructed a genomic relationship matrix with only C1 (including the PofC2). We did PCA on the kinship matrix and took the first 100 principal components as genomic predictors. We ran 1000 iterations of the genetic algorithm 10 times at each sample size. Sample sizes ranged from 200 to 2400 at increments of 400 (Supplementary Table 1). Predictions at each sample size were then made with each of 10 random and 10 optimized training sets using GBLUP in two scenarios: either just the sample of C1 was used to train the model or the sample of C1 plus all of the GG were used.

**Figure 3.**
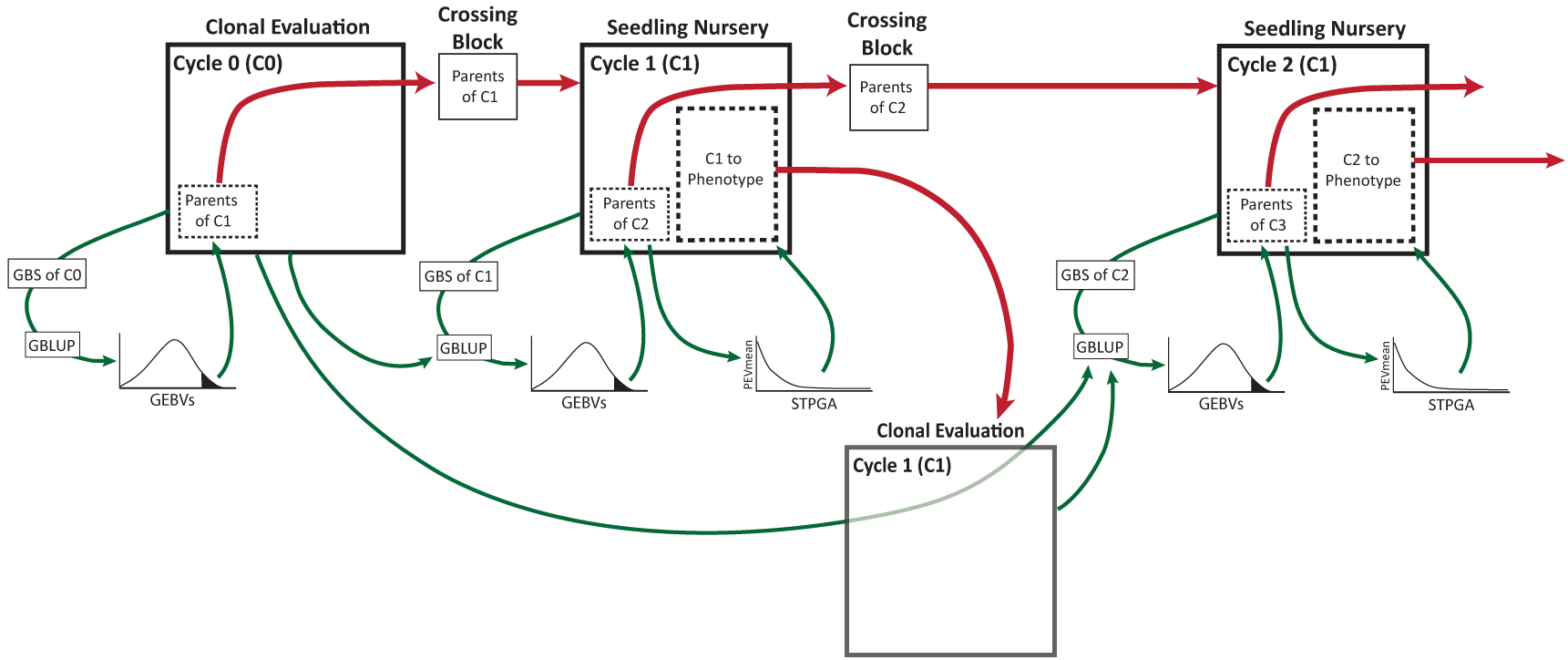
Schematic of genomic selection with training population optimization by STPGA. Selection is initially made among available, genotyped candidates based upon genomic prediction with available phenotype data. Selected parents are grown and mated in a crossing block. Resulting Cycle 1 (C1) seeds are subsequently collected and grown in a nursery. C1 seedlings are genotyped by GBS and selections are made based on genomic prediction alone. Selected parents of C2 are cloned into a crossing nursery. STPGA is used to select the optimal additional C1 seedlings to plant in a clonal evaluation trial. Because C2 seedlings do not yet exist, STPGA is instead used to select the optimal C1 seedlings to predict the selected parents of C2. Phenotypes from C1 clonal evaluation are added to the existing genomic prediction training dataset. The updated training model is used to predict breeding values of C2 seedlings when GBS data become available and the selections of parents of C3 is made. Subsequent cycles proceed based on this procedure.

### Across-population genomic predictions

We predicted all traits using all methods in three scenarios: GG (IITA Genetic Gain) +NR (NRCRI) predicts UG (NaCRRI), GG+UG predicts NR, NR+UG predicts GG (Supplementary Table 2A). Across-population predictions were made using the prediction models described above and were done following the two-step approach as also described above.

We selected optimized subsets of the combined datasets with a genetic algorithm implemented in the R package STPGA (Akdemir et al., 2015). Random subsets of the same size as the optimized subsets (300, 600, 900 and 1200) were selected for comparison between predictive accuracies. Predictions at each sample size were then made with each of 10 random and 10 optimized training sets using GBLUP.

## RESULTS

After quality control and keeping only markers with >1% MAF, the datasets had between 70,010 and 78,212 SNP markers (Table 1). Principal component analysis (PCA) of the genomic relationship matrix indicated some genetic differentiation between Nigerian populations (GG and NR) and the Ugandan training population (UG; Figure S1a). In contrast, there was little differentiation between the NRCRI and IITA GG datasets, even when comparing only the non-overlapping clones. We also calculated F_ST_ between populations as implemented in *vcftools* (Danecek et al., 2011). In agreement with results from PCA, F_ST_ between GG and NR was only 0.008, but was 0.019 and 0.021 between the Ugandan and the Nigerian populations, GG and NR, respectively. There was a similar amount of genetic differentiation between the IITA C2 progeny and its grandparental GG population (F_ST_ = 0.02) as there was between GG and UG (Table 1, Figure S1b).

**Table 1.**
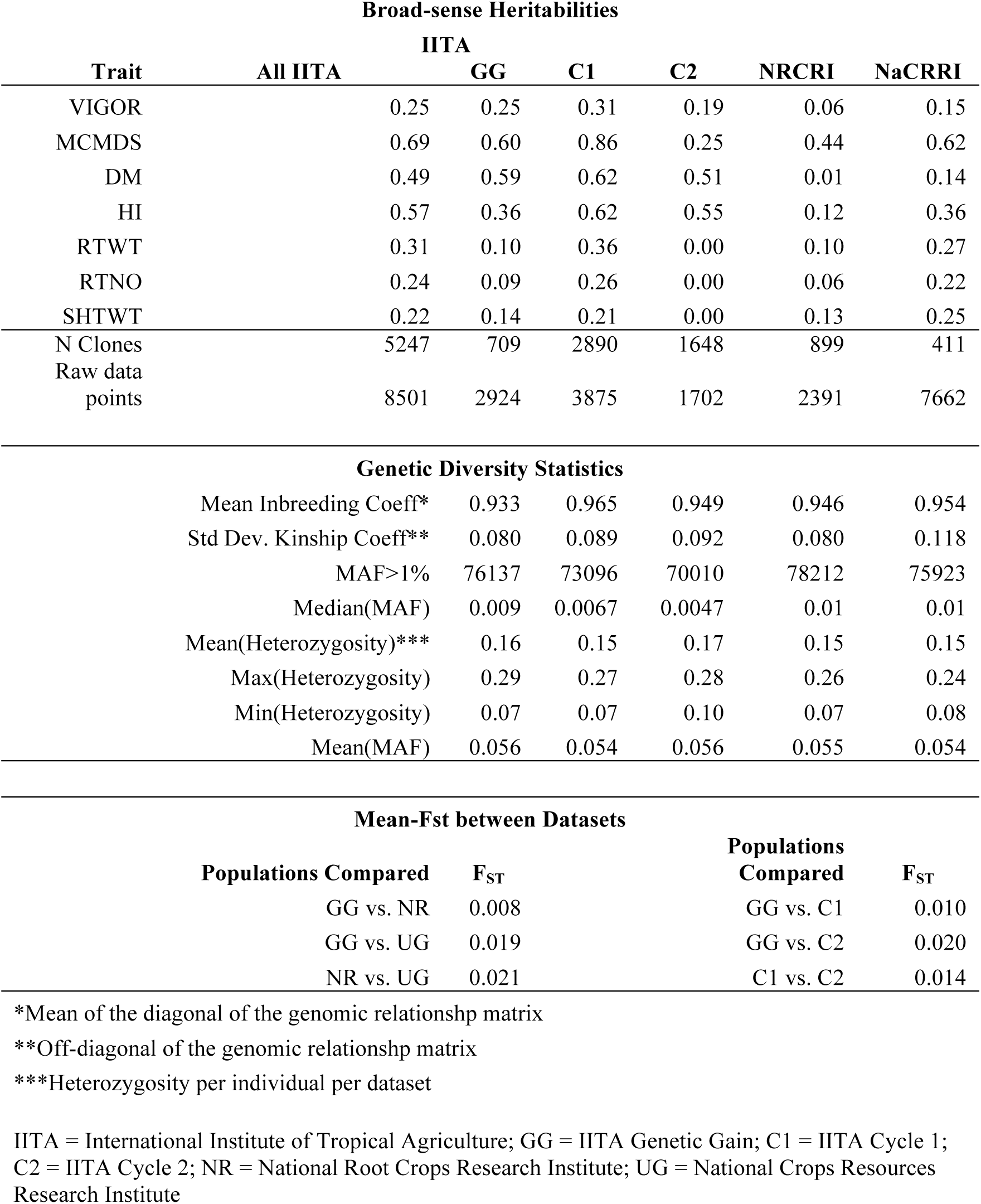
Summary and comparison of phenotype and genotype datasets analyzed in this study.

The mean inbreeding coefficient (F), as measured by the mean of the diagonal of the genomic relationship matrix, was similar for all populations, ranging from 0.933 in GG to 0.965 in C1. The mean rate of heterozygous loci was also similar between populations, ranging from 0.15 to 0.17. There was no notable decrease in heterozygosity or increase in inbreeding coefficient from GG to C1 or from C1 to C2 (Table 1; Figure S2).

In general, broad-sense heritability (H^2^) was highest in the C1 (mean 0.46 across traits), lowest for NRCRI (mean 0.13) and similar for the IITA GG and NaCRRI TPs. Averaging across populations, H^2^ was highest for MCMDS (0.57) followed by HI (0.43) and DM (0.39). However, H^2^ was generally low for yield components (Table 1).

**Table 2.**
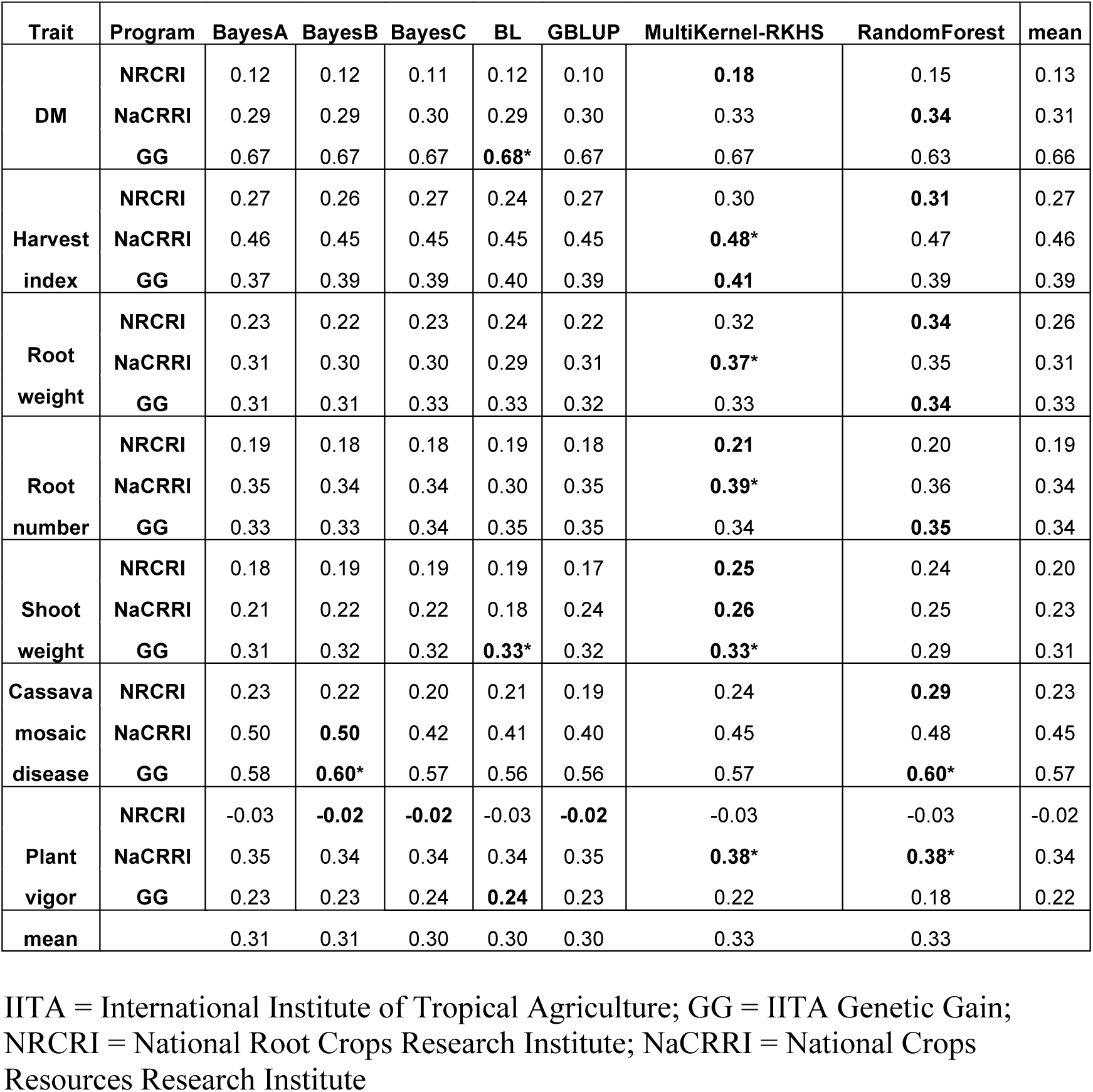
Summary of cross-validated predictive accuracies by prediction model,trait and breeding program. Highest predictive accuracy across methods within a trait and within breeding program is indicated in bold. The asterisk (*) indicates highest predictive accuracy within a trait across breeding programs

| Trait | Program | BayesA | BayesB | BayesC | BL | GBLUP | MultiKernel-RKHS | RandomForest | mean |
| --- | --- | --- | --- | --- | --- | --- | --- | --- | --- |
| DM | NRCRI | 0.12 | 0.12 | 0.11 | 0.12 | 0.10 | **0.18** | 0.15 | 0.13 |
|  | NaCRRI | 0.29 | 0.29 | 0.30 | 0.29 | 0.30 | 0.33 | **0.34** | 0.31 |
|  | GG | 0.67 | 0.67 | 0.67 | **0.68\*** | 0.67 | 0.67 | 0.63 | 0.66 |
| Harvest index | NRCRI | 0.27 | 0.26 | 0.27 | 0.24 | 0.27 | 0.30 | **0.31** | 0.27 |
|  | NaCRRI | 0.46 | 0.45 | 0.45 | 0.45 | 0.45 | **0.48\*** | 0.47 | 0.46 |
|  | GG | 0.37 | 0.39 | 0.39 | 0.40 | 0.39 | **0.41** | 0.39 | 0.39 |
| Root weight | NRCRI | 0.23 | 0.22 | 0.23 | 0.24 | 0.22 | 0.32 | **0.34** | 0.26 |
|  | NaCRRI | 0.31 | 0.30 | 0.30 | 0.29 | 0.31 | **0.37\*** | 0.35 | 0.31 |
|  | GG | 0.31 | 0.31 | 0.33 | 0.33 | 0.32 | 0.33 | **0.34** | 0.33 |
| Root number | NRCRI | 0.19 | 0.18 | 0.18 | 0.19 | 0.18 | **0.21** | 0.20 | 0.19 |
|  | NaCRRI | 0.35 | 0.34 | 0.34 | 0.30 | 0.35 | **0.39\*** | 0.36 | 0.34 |
|  | GG | 0.33 | 0.33 | 0.34 | 0.35 | 0.35 | 0.34 | **0.35** | 0.34 |
| Shoot weight | NRCRI | 0.18 | 0.19 | 0.19 | 0.19 | 0.17 | **0.25** | 0.24 | 0.20 |
|  | NaCRRI | 0.21 | 0.22 | 0.22 | 0.18 | 0.24 | **0.26** | 0.25 | 0.23 |
|  | GG | 0.31 | 0.32 | 0.32 | **0.33\*** | 0.32 | **0.33\*** | 0.29 | 0.31 |
| Cassava mosaic disease | NRCRI | 0.23 | 0.22 | 0.20 | 0.21 | 0.19 | 0.24 | **0.29** | 0.23 |
|  | NaCRRI | 0.50 | **0.50** | 0.42 | 0.41 | 0.40 | 0.45 | 0.48 | 0.45 |
|  | GG | 0.58 | **0.60\*** | 0.57 | 0.56 | 0.56 | 0.57 | **0.60\*** | 0.57 |
| Plant vigor | NRCRI | -0.03 | **-0.02** | **-0.02** | -0.03 | **-0.02** | -0.03 | -0.03 | -0.02 |
|  | NaCRRI | 0.35 | 0.34 | 0.34 | 0.34 | 0.35 | **0.38\*** | **0.38\*** | 0.34 |
|  | GG | 0.23 | 0.23 | 0.24 | **0.24** | 0.23 | 0.22 | 0.18 | 0.22 |
| mean |  | 0.31 | 0.31 | 0.30 | 0.30 | 0.30 | 0.33 | 0.33 |  |

IITA = International Institute of Tropical Agriculture; GG = IITA Genetic Gain; NRCRI = National Root Crops Research Institute; NaCRRI = National Crops Resources Research Institute

### Prediction within breeding populations

We tested seven genomic prediction models that differ by the extent and the kind of shrinkage, which is relevant to model different genetic architectures, and by their ability to capture non-additive effects (Figures S3-5).

Overall, breeding populations exhibited differences in the cross-validated prediction accuracies between methods and across traits. For NRCRI (n = 899), the mean predictive accuracy values across methods ranged between -0.02 for plant vigor and 0.27 for HI. For NaCRRI (n = 411), the mean predictive accuracy values ranged between 0.23 for shoot weight and 0.46 for HI. Meanwhile, the predictive accuracy values for GG (n = 709) ranged between 0.22 for plant vigor and 0.66 for DM.

In the NRCRI population, methods that capture non-additive effects like RKHS and random forest had the highest predictive accuracy values for all traits, except plant vigor. The trait with the highest predictive accuracy was root weight (Random forest (0.34)) and the lowest predictive accuracy was found for vigor (MultiKernel RKHS (-0.03)).

In the NaCRRI population, RKHS Multikernel showed highest predictive accuracies for all traits except for CMD, for which BayesB showed the highest value r = 0.50. In this population CMD had the overall highest predictive accuracy across traits while shoot weight exhibited the lowest predictive accuracy (Bayesian LASSO, r =0.18).

In the IITA GG population, Bayesian approaches performed better for vigor, CMD, shoot weight and DM, while RKHS method showed higher predictive accuracies for HI and for yield related traits such as root number and root weight. Meanwhile, RF gave a better predictive accuracy when used to estimate GEBVs.

Some trait-dataset combinations exhibited better predictive accuracies than others. For example, NaCRRI population had better predictive accuracies for yield components like HI, root weight and root number while the highest predictive values for CMD and DM were obtained in the GG population.

Similar to Heslot et al. (2012), we compared the cross-validated GEBV following a clustering approach. Results in Figure S6 show the hierarchical cluster trees from the combined results of the three breeding populations. Differences in clustering of methods are observed across datasets (Figure 4). In the NRCRI data, we found two groups of clustering GS methods. With BayesB, BayesC and GBLUP in one group and the rest on the other group. In the NaCRRI and IITA populations, non-parametric methods such as RKHS and Random Forest clustered together as well as the BayesA with Bayesian LASSO and GBLUP cluster with BayesC or BayesB.

**Figure 4.**
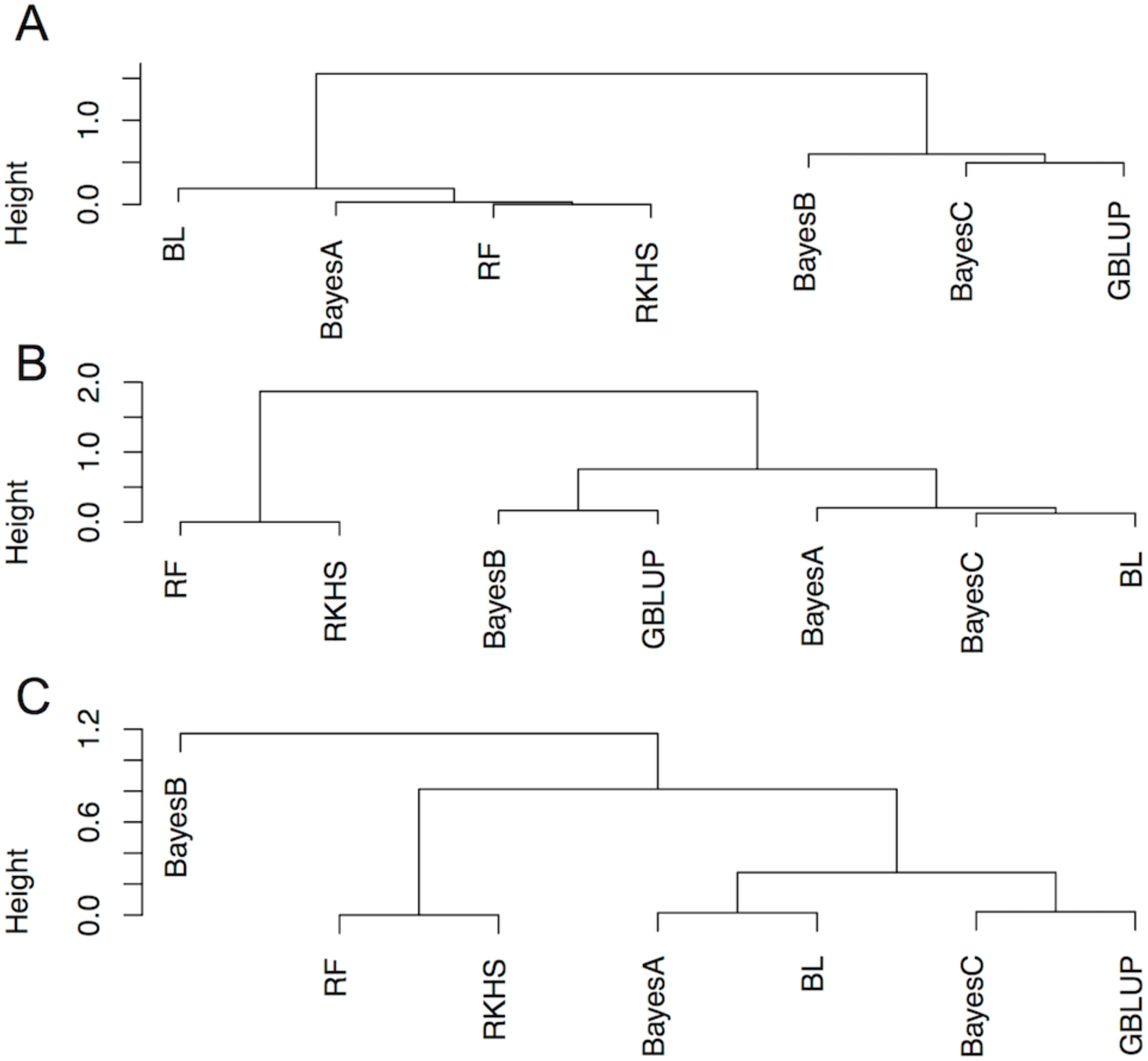
Hierarchical clustering of genomic prediction models based on cross-validated genomic estimated breeding values (GEBVs). Height on the y-axis refers to the value of the dissimilarity criterion. (A) Clustering of prediction models in the NRCRI population. (B) Clustering of prediction models in the NaCRRI population. (C) Clustering of prediction models in Genetic Gain (GG) population. GBLUP, genomic best linear unbiased predictor; BL, Bayesian Lasso; RF, random forest; RKHS, reproducing kernel Hilbert spaces multi-kernel model.

### Across-population prediction

Previous studies have reported close relatedness between the clones in the NextGen training populations (Wolfe et al., 2016). One important question within this project is whether or not datasets from different breeding programs can be combined in a training set to increase predictive accuracy. The application of any prediction model with the combined dataset would then benefit from an increase in the training population size with an outlook of using such models by other cassava breeding programs in Africa. With that in mind, we used combined datasets of GG+NR, GG+UG and UG+NR to predict the population that was not included in the training set UG, NR and GG respectively.

When predicting the traits in the UG dataset, with the combined GG+NR full set, Bayesian models gave better predictive accuracies for MCMDS, RTNO and DM. Random Forest gave better predictive accuracies for HI and RKHS for root weight and shoot weight (Table S2a).

The average predictive accuracy with the combined GG+NR full set as training set using the GBLUP model was consistently lower for all the traits when compared to the average GBLUP cross validation results (Table S2a). Furthermore, the subsets selected by STPGA to predict the NaCRRI (UG) validation set gave, for all traits and all subset sizes, lower predictive accuracies than the GBLUP cross-validation model (Table 3; Figure S7; Table S2b).

**Table 3.**
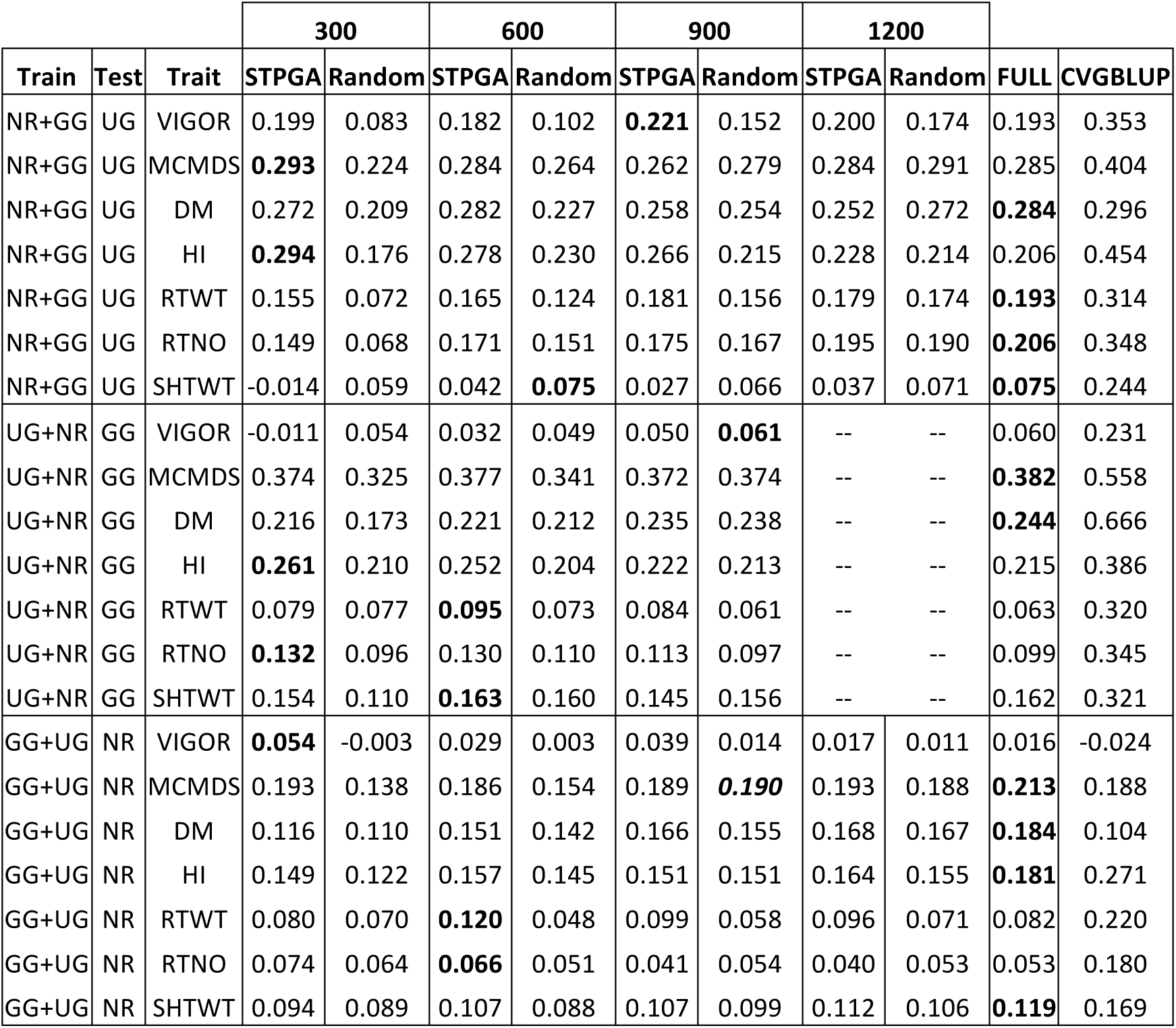
Summary of mean GBLUP cross-validated predictive accuracies cross populations. Four subset selection methods (random vs. STPGA) and the full set were considered. Highest predictive accuracy across subsets and the full set is indicated in bold, CVGBLUP:crossvalidation GBLUP within the test population. NR:NRCRI,UG:NaCRRI,GG: Genetic gain IITA.

For plant vigor, MCMDS and HI, the optimized STPGA subsets gave higher predictive accuracies than the combined GG+NR full training dataset. With few exceptions (MCMDS, SHTWT and DM) the optimized STPGA datasets gave better prediction accuracies than the same size random sets. As the optimized STPGA dataset increased in size, the predictive accuracy did not increase, except for root number where the highest predictive accuracy was found when the training population size was 1200.

When combined GG+UG full training dataset was used to predict the NRCRI training population, Random Forest and RKHS prediction models performed better for root weight, shoot weight, root number and plant vigor. Bayesian models gave better predictive accuracies for MCMDS and DM. For plant vigor, MCMDS and DM, the combined UG+GG full dataset gave better predictive accuracies than the GBLUP cross validation model (Figure S8; Table S2b). For prediction of the NRCRI training population, the optimized STPGA selected datasets gave better predictive accuracies for plant vigor, root weight, root number and shoot weight than the combined UG+ GG full training dataset.

To predict the NRCRI training population for all traits except root number (at n=900 and n=1200) and CMD (n=900), the optimized datasets gave higher predictive accuracies than the random datasets. For plant vigor, CMD resistance and DM the selection of optimized datasets with STPGA gave better predictive accuracies than the GBLUP cross validation model.

Among the STPGA datasets, the highest predictive accuracy was not always the result of an increase in training population size. For CMD resistance, the highest predictive accuracy was found, with the same value than the highest optimized size, for the smallest optimized dataset.

Predictive accuracy results of traits in the GG dataset using the full training set (UG+NR) varied across methods. Whereas Bayesian methods gave better predictive accuracy values for MCMD and plant vigor, RKHS performed better for DM, HI, root number and shoot weight. The combined (UG+NR) full training dataset for prediction of the GG population gave lower predictive accuracies than the GBLUP cross-validation model for all the traits. GBLUP cross-validation model also gave better predictive accuracies for all the traits than the random and optimized STPGA datasets. The optimized STPGA datasets gave better predictive accuracies compared to the random sets for all the traits except for plant vigor and for DM (optimized dataset n = 900) (Figure S9; Table S2b). For all traits except MCMDS and DM, the optimized STPGA subsets gave higher predictive accuracies than the combined UG+NR full training dataset.

For all the cross population results, we tested if the optimized STPGA sets would do better than random with a binomial test, assuming independence of the comparisons. We compared how many times the prediction accuracy of STPGA was greater than random for all traits. We found that for the prediction of the NR and UG sets, the STPGA optimized sets perform better than the random sets. On the contrary, when applying the same comparison of the STPGA sets with the prediction with full sets, the latter had significantly higher number of full set greater than STPGA predictive accuracy results.

Additionally, we tested if there was differential enrichment in the optimized STPGA training set of any of the populations relative to the source sets. We found a significant enrichment of the GG population (p<0.001) in the STPGA of different sizes, for the prediction of NR set using GG+UG. Similarly, we found a significant enrichment of the NR population (p<0.001), in the STPGA of different sizes, for the prediction of the GG set using the UG-NR. On the contrary, we found no significant enrichment of any population in the STPGA optimized sets for the prediction of the UG population.

### Across-generation prediction

One major area where analysis was needed concerned prediction across generations. Selections can be done at the seedling stage if GEBV can be predicted based on the previous generations and training data. Because nearly all of the IITA germplasm from C1 and C2 were clonally evaluated, we were able to use these data to assess the accuracy of genomic prediction on unevaluated genotypes of the next generation. In general, the accuracy of prediction across generation was greatest when predicting C2 as evidenced by averaging across prediction models and traits for predictions trained either with C1 (mean 0.19 ± standard error 0.02) or GG+C1 (0.19 ± 0.02). The accuracy was lower on average when predicting C2 with GG (0.11 ± 0.01) compared to predicting C1 with GG (0.17 ± 0.02). Accuracy was lowest for both VIGOR and RTWT (0.06 ± 0.005) and highest for MCMDS (0.32 ± 0.03) and DM (0.38 ± 0.01). Most prediction models performed similarly as evidenced by the averaged accuracy across traits and training-test combinations with RF performing worst (0.08 ± 0.01) and BayesA and BayesB performing best (both 0.20 ± 0.03). For MCMDS, we found that prediction accuracy was greatest using BayesA and BayesB (Figure 5, Figure S10, Table S3).

**Figure 5.**
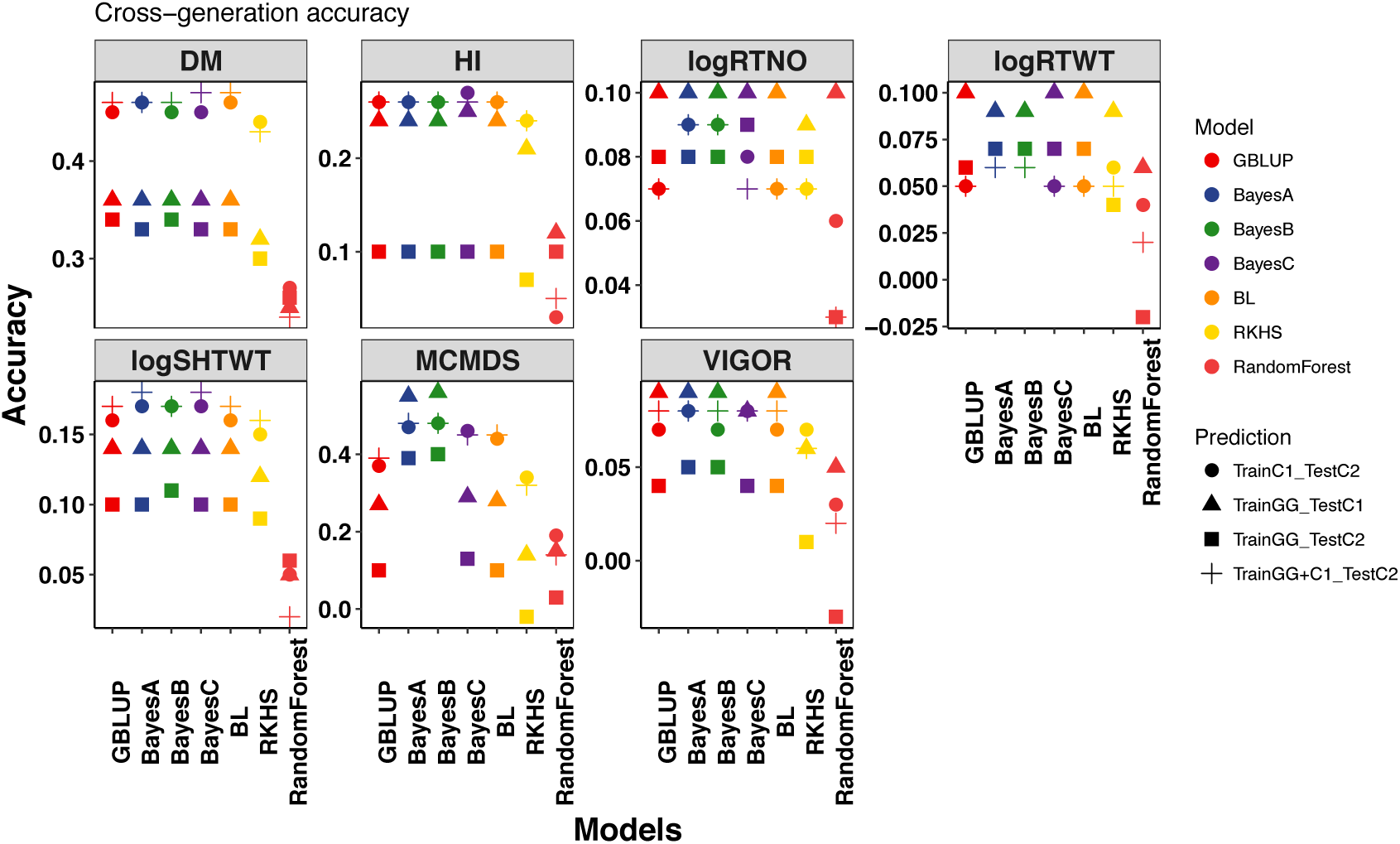
Boxplot of cross-generation prediction accuracies. Seven genomic prediction methods were tested for seven traits (panels). For each model – trait combination, four predictions were made: GG predicts C1, GG predicts C2, C1 predicts C2, GG+C1 predicts C2. Boxes show range of accuracies across these four prediction scenarios. All data are from the IITA Genomic Selection program. GG=Genetic Gain. C1 = Cycle 1. C2 = Cycle 2.

### Training population update

The first 100 PCs of the C1 kinship matrix were used as predictors for STPGA and explained 97.7% of the genetic variance. In all cases the genetic algorithm converged within the 1000 iterations run (Figure S11).

Given the constraints of breeding programs described above, it was necessary to select samples of C1 that were optimized for predicting the parents of C2 (PofC2), rather than the C2 themselves. Despite targeting the PofC2, we used selected training sets to predict C2, thus simulating the addition of phenotypes to the training set. Because of this, we compared the accuracy of subsets of C1 predicting C2 to accuracy predicting the PofC2. As the number sampled increased from 200 to 2,400, averaging across traits and methods for subset selection (STPGA and Random), accuracy increased by 120 and 105% when predicting C2 and PofC2, respectively. Accuracy increase was smaller when including the 709 GG clones in the prediction, increasing only by 43 and 36% respectively when predicting C2 and PofC2 (Supplementary Table 4).

STPGA consistently selected training datasets with lower expected mean PEV on the test set compared to random and across training set sizes (Figure S12). Further, using STPGA to select clones for phenotyping gave an average 13% better accuracy (average accuracy of 0.242 vs. 0.214, two-tailed t=6.29, df=4458, p<0.0001) compared to random sampling. Broken down by validation set, STPGA was significantly better than random predicting PofC2 (t=9.8, df=2147, p<0.0001), but not significantly better for predicting C2 (t=1.41, df=2227, p=0.16).

We compared these accuracies with that of the full set of C1 (or GG+C1) *and* to the cross-validation accuracy within the test set (C1 for prediction of PofC2, C2 for predictions of C2). When predicting C2, which was our primary goal, subsets were almost always inferior to the full set, with the exceptions of the middle sizes for RTWT, but the advantage was very small (Figure 6, Figure S13). However, STPGA-selected subsets tended to have better accuracy than the full set, especially for yield components when predicting the PofC2, which were the genotypes targeted by the optimization algorithm (Figure 7, Figure S14).

**Figure 6.**
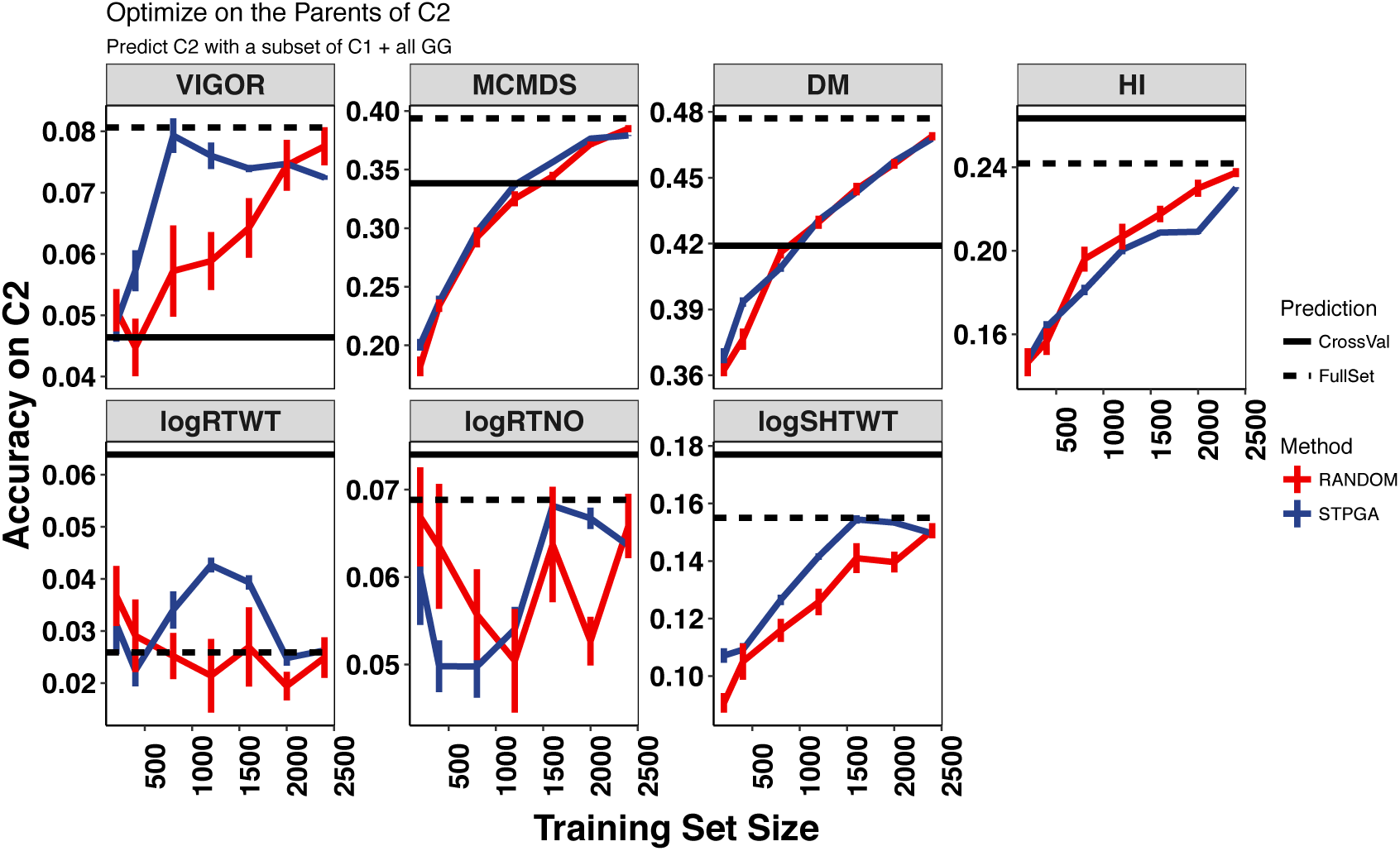
The relationship between training set size and accuracy predicting IITA Cycle 2 (across-generation). The accuracy of prediction for seven traits (panels) with the IITA Genetic Gain (GG) population training data plus data from different size subsets (x-axis) of their progeny, Cycle 1 (C1) is shown. Subsets of a given size were selected either at random or using the genetic algorithm implemented in the R package STPGA. Ten random and ten STPGA-selected s ubsets were made at each training set size. Error bars are the standard error around the mean for the ten samples. Horizontal black lines show the mean cross-validation accuracy for the C2 (validation set; solid line) and the accuracy of the full set of GG+C1 predicting C2 (dashed line).

**Figure 7.**
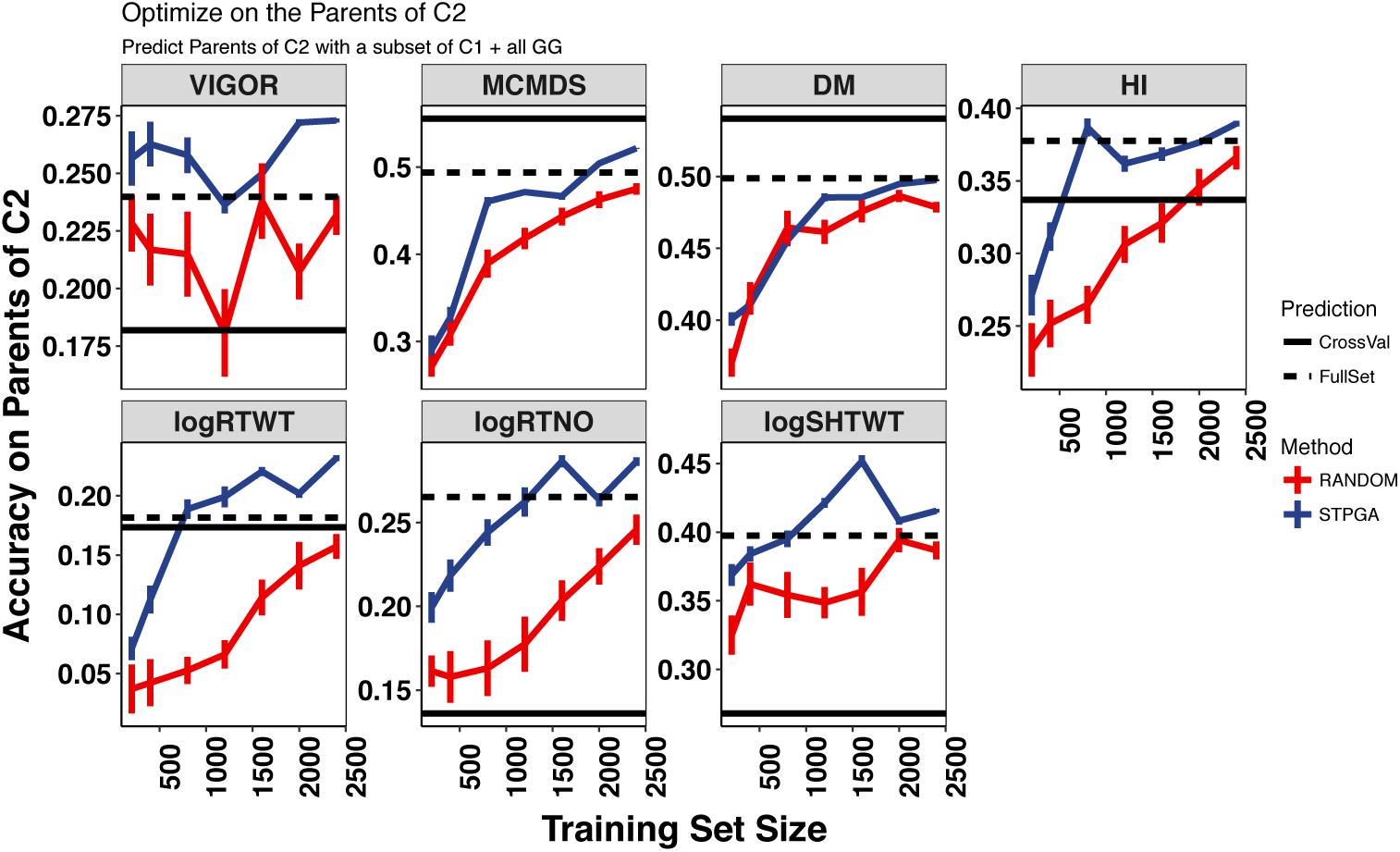
The relationship between training set size and accuracy predicting the parents of Cycle 2 (from Cycle 1, within-generation). The accuracy of prediction for seven traits (panels) with the IITA Genetic Gain (GG) population training data plus data from different size subsets (x-axis) of their progeny, Cycle 1 (C1) is shown. Subsets of a given size were selected either at random or using the genetic algorithm implemented in the R package STPGA. Ten random and ten STPGA-selected subsets were made at each training set size. Error bars are the standard error around the mean for the ten samples. Horizontal black lines show the mean cross-validation accuracy for the C1 (validation set; solid line) and the accuracy of the full set of GG+C1 predicting the parents of C2 (dashed line).

The correlation between the selection criterion, PEVmean, used by STPGA and the training set size is strong for all traits (range -0.57 to -0.61). Aside from simply increasing the TP size, we wanted to assess the extent to which the PEVmean could be used as a predictor of the achievable accuracy. Regression of prediction accuracies for each sample (regardless of whether it was selected randomly or by STPGA) on PEVmean explains between 8% (RTNO) and 46% (DM) of the variance in accuracy. Multiple regression including PEVmean and training set size (Ntrain) as predictors showed PEV to be the more significant predictor (across all traits). In fact, Ntrain was not a significant explanatory variable for RTWT or RTNO (Table S5).

## DISCUSSION

The Next Generation Cassava Breeding Project (www.nextgencassava.org) aims to assess the potential of genomic selection in cassava to reduce the length of the breeding cycle and increase the number of crosses and selection per unit time. The project is implementing genomic selection in three breeding programs from Nigeria and Uganda, with genotypic and phenotypic data from training populations and two cycles of selection available on a database dedicated to cassava (www.cassavabase.org).

Using a cross-validation scheme, we contrasted the performance of GBLUP, RKHS (Single-kernel and Multi-kernel), BayesA, BayesB, BayesCpi, Bayesian LASSO and Random Forest for yield components (RTWT, RTNO, SHTWT, HI, DM) and CMD resistance data from the breeding programs.

In general, the performance of predictive models is known to be conditional on the genetic architecture of the trait under consideration (Daetwyler et al., 2010; Su et al., 2014). While non-additive models including RF and RKHS capture dominance and epistasis effects, GBLUP is more suitable for prediction when traits are determined by an infinite number of unlinked and non-epistatic loci, with small effect.

Not surprisingly, heritability varied between populations, conceivably as a consequence of the differences in the number and design of field trials between breeding programs. For most traits, it is not possible to determine exactly the reason for differences in heritability. However, for DM, we can hypothesize that differences in phenotyping protocols between programs (specific gravity method at NRCRI and NaCRRI versus oven drying at IITA) could account for differences We note the estimate of zero heritability for RTWT, RTNO and SHTWT in the IITA C2 and acknowledge this is likely to account for the quality of cross-generation prediction of that dataset.

Cross-validation results were mostly consistent across breeding programs and the superiority of one prediction method over the others was trait-dependent. RF and RKHS usually predicted phenotypes more accurately for yield-related traits, which are known to have a significant amount of non-additive genetic variation (Wolfe et al 2016b). Similar findings have been made in wheat, for grain yield, an additive and epistatic trait, in which RKHS, radial basis function neural networks (RBFNN), and Bayesian regularized neural networks (BRNN) models clearly had a better predictive ability than additive models like BL, Bayesian ridge-regression, BayesA, and BayesB (Perez-Rodriguez et al., 2013).

While cross validation results within breeding programs are encouraging for the use of genomic selection, across breeding program prediction values were fairly low. Mean F_ST_ values lower than 0.05 indicated that the three breeding populations share genetic material. Despite this, our results indicate that the prospect for sharing data across Africa to assist in genomic selection is limited to certain traits (most notably MCMDS) and populations. Indeed, obtaining a larger training set by combining training population did not always lead to higher prediction accuracies compared to what could already be achieved within that population as evidenced by cross-validation.

In animal models, prediction with multi-breed populations has also been shown to be poor with most of the observed accuracy due to population structure (Daetwyler et al., 2012). An alternative kernel function has been proposed to estimate the covariance between individuals based on markers, which can improve fit to the data to account for genetic heterogeneity of breeding populations (Heslot and Jannink, 2015).

Conceivably, in our study the addition of individuals from different breeding programs was detrimental due to the inconsistent heritability for most traits. Another possibility is genotype-by-environment (GxE) interaction. The impact of GxE interaction on predictive accuracy has been reported in wheat when the same population was evaluated in different environments (Crossa et al., 2010; Endelman, 2011). Similarly, in cassava using historical data from the IITA’s GG population, prediction across locations led to a decrease in accuracy (Ly et al., 2013).

Using the training sets selected based on optimized algorithm gave better predictive ability than randomly assigned samples with a decrease in accuracy when compared with GBLUP cross-validation results. Although in previous studies predictive accuracies with full sets were lower than optimized subsets (Rutkoski et al., 2015), in our study we found the contrary, indicating that a larger training set was more advantageous. Combining data from different experiments and populations for across population prediction remains promising for traits like CMD where GWAS results indicate a stable large-effect QTL throughout the tested breeding populations (Wolfe et al., 2016).

When predicting unevaluated progenies from the next generation (cross generation), our results indicated, in our judgment, that accuracy should is sufficient for DM, MCMDS and to a lesser extent HI. Although accuracy is stable across the generations tested for DM using most models, for MCMDS to be successful, we recommend using a Bayesian shrinkage model such as BayesA or BayesB. The advantage of these models for CMD resistance over GBLUP likely comes because of the major known QTL segregating in the population (Rabbi et al., 2014; Wolfe et al., 2016a) and the ability of these two models to allow differential contribution of markers near the QTL to the prediction. One disadvantage of BayesB, in particular, is that the known polygenic background resistance for CMD may become dee-mphasized, in favor of heavy selection on the major effect gene(s) (Hahn et al., 1980; Legg and Thresh, 2000; Akano et al., 2002; Rabbi et al., 2014; Wolfe et al., 2016).

We noted that RF and RKHS performed poorly across generations; this is a result that makes sense given that the predictability of epistatic and dominant interactions declines with recombination (Lynch and Walsh, 1998).

Based on the datasets analyzed in this study, it was apparent that the size of a training population had a significant impact on prediction accuracy for most traits. Thus, breeding programs will benefit from phenotyping the maximum possible amount. In agreement with the results in other crops (Rincent et al., 2012; Akdemir et al., 2015; Isidro et al., 2015), our results do indicate that optimization algorithms like STPGA can provide at least a small advantage over random selections of materials for phenotyping.

Each breeding program will need to determine the amount of phenotyping vs. genotyping to do in order to maximize prediction accuracy and selection gain based on the cost and availability of land, labor and genotyping. An analysis in barley by Endelman et al. (2014) provides a good example of the potential complexity of these decisions. The authors show, as we do, that larger number of phenotyped individuals is always beneficial, and that it is usually beneficial to focus on evaluating new lines at the expense of additional phenotyping of old lines. However, if genotyping costs are high, the cost-benefit balance shifts towards more evaluation of existing lines (Endelman et al., 2014). Endelman et al.’s (2014) study focused on prediction in biparental populations. Although this is likely to apply to cassava breeding populations, we stress the necessity of doing such an analysis for each breeding application separately.

An important result is that STPGA was able to find subsets that were better than the full set for predicting the parents of C2 (PofC2). PofC2 are members of C1 and were the individuals targeted with STPGA. One possible interpretation is that the benefit comes from phenotyping contemporaries. If that were true, we could make a significant difference in accuracy by phenotyping a subset of clones from the current generation before predicting GEBV for the entire set of selection candidates. To do this without lengthening the selection and recombination cycle, harvested stems would need to be stored long enough for phenotypic data to be curated, predictions and selections to be conducted and STPGA to be run. Methods to store cassava stakes for up to 30 days are available, indicating such a scheme could be possible (Sungthongw et al., 2016). Even without improved stem cutting storage, this could be done while only lengthening the selection and recombination cycle to perhaps 1.5-2 years, which would still be significantly faster than conventional cassava breeding.

A related possibility is to place annual selection pressure on traits that are predictable across generation (e.g. MCMDS, HI and DM). Predictions of total genetic value for yield traits for selection of clones that will be tested as potential varieties could then be done after clonal evaluation data become available on at least a subset of contemporary genotypes. Further trials will be necessary to determine whether there is an advantage to this type of strategy.

The primary promise genomic selection offers to cassava breeding is the ability to select and recombine germplasm more frequently and thus hopefully speed the rate of population improvement while combining a myriad of quality, disease and yield related traits into a single genotype that can be released as a variety. The applicability of results from the different prediction models in cassava is then dependent on whether the goal is the prediction of breeding value of progeny or the selection of advanced lines for testing as varieties.

We are still in the early stages of GS in this crop, but results are promising, at least for some traits. The TPs need to continue to grow and quality phenotyping is more critical than ever. However, general guidelines for successful GS are emerging. Phenotyping can be done on fewer individuals, cleverly selected, making for trials that are more focused on the quality of the data collected.

## ACKNOWELDGEMENTS

We acknowledge the Bill & Melinda Gates Foundation and UKaid (Grant 1048542; http://www.gatesfoundation.org) and support from the CGIAR Research Program on Roots, Tubers and Bananas (http://www.rtb.cgiar.org). We give special thanks to A. G. O. Dixon for his development of many of the breeding lines and historical data we analyzed. Thanks also to A. I. Smith and technical teams at IITA, NRCRI and NaCRRI for collection of phenotypic data and to A. Agbona, P. Peteti, A. Ogbonna, E. Uba and R. Mukisa for data curation.

### CONFLICTS OF INTEREST

No conflicts.

